# Diversity-enhanced canopy space occupation and leaf functional diversity jointly promote overyielding in tropical tree communities

**DOI:** 10.1101/2024.06.06.597699

**Authors:** Tama Ray, Andreas Fichtner, Matthias Kunz, Tobias Proß, Pia M. Bradler, Helge Bruelheide, Louis Georgi, Sylvia Haider, Michaela Hildebrand, Catherine Potvin, Florian Schnabel, Goddert von Oheimb

**Affiliations:** Institute of General Ecology and Environmental Protection, Technische Universität Dresden, Tharandt, Germany; German Centre for Integrative Biodiversity Research (iDiv) Halle-Jena-Leipzig, Leipzig, Germany; Institute of Biology/Geobotany and Botanical Garden, Martin Luther University Halle-Wittenberg, Halle (Saale), Germany; Institute of Ecology, Leuphana University of Lüneburg, Lüneburg, Germany; Helmholtz Centre Potsdam - GFZ German Research Centre for Geosciences, Telegrafenberg, 14473 Potsdam, Germany; ThüringenForst-AöR State Forest, Hallesche Straße 20, 99085 Erfurt, Germany; Chair of Silviculture, Institute of Forest Sciences, University of Freiburg, Tennenbacherstr. 4, 79085, Freiburg, Germany; Department of Biology, McGill University, 1205 Dr Penfield, Montréal, Québec, Canada, H3A 1B1; Smithsonian Tropical Research Institute, Panama, Panama

## Abstract

Understanding the mechanisms that drive biodiversity-productivity relationships is critical for guiding forest restoration. Although complementarity among trees in the canopy space has been suggested as a key mechanism for greater productivity in mixed-species tree communities, empirical evidence remains limited. Here, we used data from a tropical tree diversity experiment to disentangle the effects of tree species richness and community functional characteristics (community-weighted mean and functional diversity of leaf traits) on canopy space filling, and how these effects are related to overyielding. We found that canopy space filling was largely explained by species identity effects rather than tree diversity effects. Communities with a high abundance of species with conservative leaf traits were those with most densely packed canopies. Overall, a higher canopy space filling translated into an enhanced wood productivity, with communities associated with a high taxonomic and functional diversity being the most productive. Importantly, most communities (83%) produced more wood volume than the average of their constituent species in monoculture (i.e. most communities overyielded). Our results show that overyielding increased with leaf functional diversity and positive net biodiversity effects on canopy space filling, which mainly arose due to a high taxonomic diversity. These findings suggest that both taxonomic diversity-enhanced canopy space filling and canopy leaf diversity are important drivers for overyielding in mixed-species forests. Consequently, restoration initiatives should promote stands with functionally diverse canopies by selecting tree species with large interspecific differences in leaf nutrition, as well as leaf and branch morphology to optimize carbon capture in young forest stands.

## 1. Introduction

Forest restoration is gaining importance due to the twin crises of climate change and species extinction, following a long period of forest loss caused by deforestation, land-use change, or degradation. This is emphasized by nature-based solution initiatives such as the UN Decade of Ecosystem Restoration and the Bonn Challenge (FAO, SER and IUCN CEM, 2023). Forest restoration plays a crucial role in providing multiple ecosystem functions and services such as carbon sequestration, climate regulation, and energy and water supply, while also helping to reduce biodiversity loss (Brancalion et al., 2019; Diaz et al., 2006; Koch and Kaplan, 2022). Recent experimental and observational studies have increasingly demonstrated a positive relationship between the provision of forest ecosystem functions and services and biodiversity (Gamfeldt et al., 2013; Huang et al., 2018; Liang et al., 2016; Veryard et al., 2023; Gao et al., 2021). Specifically, the planting of mixed-species forests (Messier et al., 2021; Puettmann et al., 2009) can achieve higher biomass productivity in restoration projects, as diverse plantations have been shown to be more productive than monocultures (Depauw et al., 2024; Fichtner et al. 2018). The mechanisms that drive these biodiversity-productivity relationships (BPRs) in forests are, however, still under debate.

In a recent study of a young experimental tree plantation in temperate Europe, Ray et al. (2023) found that tree species richness increased wood productivity by enhancing the aboveground stand structural complexity. This supports the niche complementarity hypothesis, which proposes that increasing species richness improves resource-use efficiency and enhances ecosystem functioning (Tilman, 1997). Although other studies have also shown the positive impact of tree diversity on structural complexity within a stand (Juchheim et al., 2019; Zemp et al., 2019; Perles-Garcia et al., 2021; Coverdale and Davies, 2023), the effects of structural complexity or structural diversity on stand productivity have yielded equivocal results (Dănescu et al., 2016; Schnabel et al., 2019). Juchheim et al. (2017) propose that these inconsistent findings may be attributed to measures that fail to encompass those structural attributes that significantly impact stand growth, such as the spatial arrangement of tree crowns in the canopy. The crown of a tree is crucial for its productivity because leaves’ position and distribution largely determine the absorption of photosynthetically active radiation or the light-use efficiency (Niinemets, 2010; Valladares, 1999). Many tree species have a high crown plasticity, allowing them to improve their light absorption by growing their branches towards areas with higher light intensity (Fichtner et al., 2013; Perles-Garcia et al., 2022; Pretzsch, 2014). Consequently, tree species mixing enables greater spatial complementarity among trees in the canopy space due to vertical stratification (Ishii and Asano, 2010) and diversity-induced crown plasticity (Kunz et al., 2019; Lang et al., 2010; Pretzsch, 2014; Williams et al., 2017; Ali et al., 2019), which is generally referred to as crown complementarity. A greater crown complementarity in turn, often results in more densely packed canopies (Jucker et al., 2015). Therefore, it is crucial to quantify canopy space filling (also known as canopy packing or canopy space occupation; Georgi et al., 2022; Jucker et al., 2015; Pretzsch, 2014; Seidel et al., 2013) to deepen our understanding on the relationship between structural diversity and stand productivity (Juchheim et al., 2017).

Canopy space filling is calculated as the proportion of the space potentially available for tree crowns that are already occupied by the crown volumes of all existing trees within a stand (Pretzsch 2014). In the past, the analysis of canopy space filling has primarily involved rough estimates of the volume of individual tree crowns using basic geometric shapes derived from manual field measurements of tree height, crown radii, and crown depth (e.g. Jucker et al., 2015; Lang et al., 2012; Martin-Blangy et al., 2023; Pretzsch, 2014). There is however a significant limitation to this approach: the crown volume alone may not be sufficient to test for diversity effects in mixtures, as individual-tree crowns can experience significant diversity-induced changes in leaf area or branching density (Guillemot et al., 2020; Hildebrand et al., 2021; Kunz et al., 2019; Sapijanskas et al., 2014). Therefore, detailed considerations of the individual crowns are essential to accurately assess tree diversity effects on canopy space filling in mixtures.

Terrestrial laser scanning (TLS) is a modern technology that can capture the crowns of individual trees with great accuracy and high efficiency (Calders et al., 2020; Seidel et al., 2011; Hess et al., 2018; Lines et al., 2022). For the precise quantification of stand-level canopy space filling, the three-dimensional TLS point clouds are converted into a grid of voxels (i.e. cubes with a defined edge length) and the voxels containing laser points of the tree crowns (i.e. “filled voxels”) are identified (Seidel et al., 2013). It is important to note that the individual tree crowns are directly integrated with their entire structure and there is no need to approximate the crown with a simple geometric shape (Juchheim et al., 2017; Hess et al., 2018). However, TLS-based quantifications of canopy space filling have rarely been applied in tree diversity research so far (but see Georgi et al., 2022; Seidel et al., 2013). Specifically, we lack experimental evidence of canopy space filling-productivity relationships in controlled tree biodiversity–ecosystem functioning (BEF) experiments.

The functional composition of a community, which is determined by the functional traits of its constituent species, can also shape biodiversity-productivity relationships (BPRs). Two main trait-based mechanisms can explain positive BPRs: First, the functional identity of the dominant species drives community productivity, which is consistent with the mass-ratio hypothesis (Grime, 1998) and expressed as the community-weighted mean (CWM) of a particular trait. Second, according to the niche complementarity hypothesis (Tilman, 1997), the combination of functionally dissimilar species (i.e. communities’ functional diversity; FD) allows for a higher community productivity through resource partitioning or facilitation.

For example, in a sub-tropical forest in China found that both FD and CWM both could strongly support productivity (Ali et al., 2017), but in a subtropical tree BEF experiment, Bongers et al. (2021) found that over time FD became a stronger predictor of productivity than CWM. Leaf traits are closely related to species’ resource-use strategies (Wright et al., 2004) and productivity (Poorter and Bongers, 2006), and thus can capture the effects of communities’ functional identity and diversity on productivity. For example, a plant’s ability to acquire light and conduct photosynthesis is determined by leaf traits such as specific leaf area and leaf nitrogen concentration (Milla and Reich, 2007; Reich et al., 1998). In contrast, leaf dry matter content, leaf carbon content, and the ratio of leaf carbon to nitrogen are associated with leaf longevity and structural integrity (Niinemets et al., 2014). This trade-off between increased photosynthetic rate and leaf construction (Onoda et al., 2017) is captured by the leaf economics spectrum (Wright et al., 2004), which describes differences in resource-use strategies among species: rapid resource acquisition (acquisitive species) versus greater resource conservation (conservative species). Thus, a higher canopy space filling and productivity can either emerge through an optimization of leaf physiology (Niinemets, 2012), which should be reflected in changes of CWM of a specific leaf trait, or through an increase in leaf functional diversity (Plekhanova et al., 2021). However, it remains unclear whether the dominance of a tree species with a particular leaf trait in a community (CWM) or a larger leaf trait diversity (FD) drives canopy space filling, and how tree diversity-induced changes in canopy space filling affect community productivity.

To improve our understanding of diversity-induced changes in canopy space filling and its link to community productivity, we used data from the oldest tropical forest BEF experiments, the Sardinilla experiment, Panama (Potvin and Dutilleul, 2009; Scherer-Lorenzen et al., 2005). Being located in the tropics, this experiment can therefore provide valuable insights for forest restoration in this biome (Brancalion et al., 2019). Previous studies in the Sardinilla experiment provided evidence for positive effects of tree species diversity and structural diversity on productivity (Potvin and Gotelli, 2008; Schnabel et al., 2019; Guillemot et al., 2020), but the underlying mechanisms remained unclear. In this study, we therefore used high-resolution TLS point cloud data to assess the three-dimensional canopy structure combined with leaf trait measurements that account for intraspecific trait variation to identify processes that drive changes in canopy space filling along a tree species richness gradient (ranging from monocultures to five-species mixtures) and how these changes are linked to community productivity. We conducted a detailed analysis of the canopy space, including potential variations in individual-tree crown architecture in response to neighborhood diversity, which has been previously reported from the Sardinilla experiment (Guillemot et al., 2020) and other BEF experiments (e.g. Kunz et al., 2019; Hildebrand et al., 2021). Additionally, we included data on leaf traits and tree growth measured at the Sardinilla experimental site to better understand canopy-related processes for enhanced productivity in tree species mixtures. Specifically, we (i) explored the relative importance of leaf functional community characteristics (CWM and FD) and tree species richness for canopy space filling, (ii) explored the relationship between canopy space filling and community productivity along the tree species richness gradient, and (iii) assessed the relative importance of canopy space filling, tree species richness and leaf functional community characteristics (CWM and FD) for driving overyielding in wood volume (i.e., higher wood volume in a mixture compared to the weighted average of monoculture yields).

## 2. Materials and Methods

### 2.1. Study site and design

This study was conducted in a 16-year-old tropical tree diversity experiment established in Sardinilla, 55 km north of Panama City (9° 19’N, 79° 38’W). The climate at the study site is tropical with an annual mean temperature of 26 °C and annual precipitation of 2,661 mm (BCI, Physical Monitoring Program of the Smithsonian Tropical Research Institute).

After deforestation of the native semi-deciduous lowland forest in 1952/1953, the area of 5 ha was first used for agriculture and then converted to pasture (Scherer-Lorenzen et al., 2005). A total of 5,566 <6-month old seedlings were planted at a distance of 3 m, following standard reforestation practices in Panama (Potvin and Gotelli, 2008). The experiment originally consisted of six native tree species planted in monocultures, three- and six-species mixtures (Potvin and Dutilleul, 2009): *Anacardium excelsum* (Bertero & Balb. Ex Kunth) Skeels (Anacardiaceae), *Cedrela odorata* L. (Meliaceae), *Cordia alliodora* (Ruiz. & Pav.) Oken (Boraginaceae), *Hura crepitans* L. (Euphorbiaceae), *Luehea seemannii* Triana & Planch. (Malvaceae) and *Tabebuia rosea* (Bertol.) A. DC. (Bignoniaceae). These tree species were planted in 24 plots of 45 x 45 m each: 12 plots with monocultures, six plots with different three-species mixtures, and six plots with mixtures of all six species (Potvin and Dutilleul, 2009; Scherer-Lorenzen et al., 2005). The species composition of the three-species mixtures was randomly selected based on their growth rates and the frequency of occurrence in gaps or closed forests of the permanent plot on Barro Colorado Island (BCI) to ensure trait divergence (Scherer-Lorenzen et al., 2005). Each three-species mixture consisted of a pioneer species considered to be fast-growing (either *L. seemannii* or *C. alliodora*), a light-intermediate species (either *A. excelsum* or *H. crepitans*), and a shade-tolerant species considered to be slow-growing (either *T. rosea* or *C. odorata*) (Scherer-Lorenzen et al., 2005). Plots were further divided into four subplots (quarter quadrants) and we used these subplots nested within plots in our analysis. Due to high mortality (>85%) of *C. alliodora* during the first years of the experiment (Potvin and Gotelli, 2008; Scherer-Lorenzen et al., 2007), monoculture data for this species were not available. Although *C. alliodora* was still present in 12 subplot mixtures in 2017, we excluded these subplots from the subsequent analyses to ensure unbiased comparisons between monocultures and mixtures. This resulted in a total of 21 plots and 76 subplots. Overall, the average tree mortality in these subplots was comparatively low during the study period 2012-2017 (median: 2.19%, mean: 5.13%). The realized gradient of tree species richness in the subplots therefore ranged from monocultures (n_subplots_=40) to two- (n_subplots_=7), three- (n_subplots_=10), four- (n_subplots_=2) and five-species mixtures (n_subplots_=17).

### 2.2. Terrestrial laser scanning data acquisition and post processing

TLS data collection took place in early June 2017 using a RIEGL VZ-400i (RIEGL, Austria), which provides a 360° (horizontal) x 130° (vertical) field-of-view and full-waveform analysis capabilities. Scanning was performed on dry and calm days with temperatures around 25 °C (for more details see Guillemot et al., 2020).

The plots were scanned in a multi-scan mode, resulting in 16 scan positions per plot with two scans per position: One scan was performed with the TLS scanner in a vertical position and the other in a horizontal (90° tilt) orientation to represent the entire canopy space. The scanner operates at a wavelength of 1,550 nm, and the angular scan resolution was set to 0.04°, which corresponds to a point sampling distance of approximately 7 mm at 10 m distance. In addition, the scan frequency was 600 kHz to achieve better canopy penetration of each scan pulse (Guillemot et al., 2020). To account for scan shadows, the distance between scan positions was approximately 10 m in a grid pattern to capture each tree from multiple positions to represent the 3D tree structures.

Using the RiSCAN Pro software (version 2.6.2), we co-registered the point clouds of the individual scans of each plot. The registration was performed using the multi-station adjustment approach with plane patches in the RiSCAN Pro software with a relative registration accuracy between scans within 3 mm (Guillemot et al., 2020). The point clouds were cut with lasclip (LAStools, 2022) using the coordinates of each plot. To avoid side effects from neighboring plots or open areas, a 2 m wide buffer was placed around the outer edges before cutting the four subplots per plot.

### 2.3. Canopy space filling analysis

Canopy space filling was determined for each subplot following the approach of Georgi et al. (2022). The upper boundary of the canopy space was defined as the highest tree-top of each subplot, and the lower boundary was defined as the lowest crown base height of each subplot (Figure 1a). This approach is similar to the canopy space definitions used by Seidel et al. (2013) and Jucker et al. (2015). The canopy space of each subplot was voxelized with lasvoxel (LAStools, 2022) using a voxel size of 0.4 m edge length to minimize occlusion effects of tropical trees in leaf-on conditions according to Béland et al. (2011). The relative canopy space filling is defined as the percentage of the occupied voxels out of the total voxelized canopy space (hereafter referred to as canopy space filling index, CSFI).

**Figure 1.**
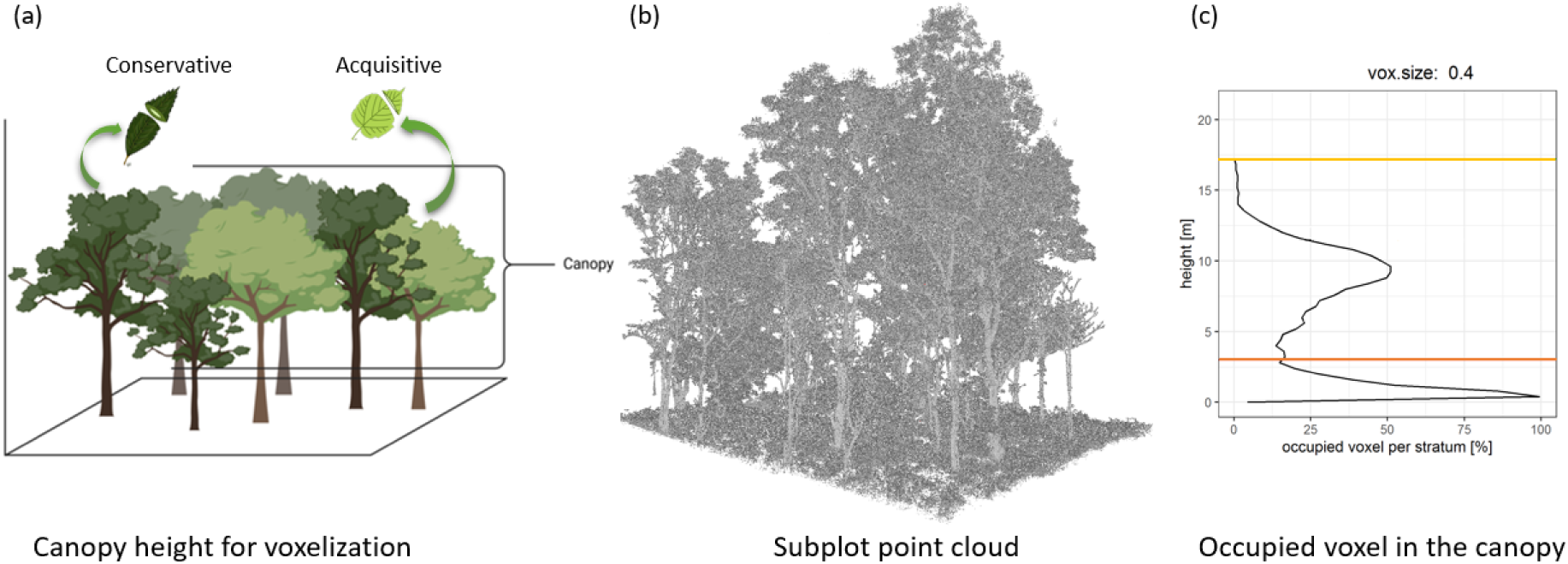
**(a)** Conceptual figure illustrating canopy space filling by species with conservative and acquisitive leaf traits and growth strategies, respectively. A greater leaf trait diversity is assumed to result in more complementary canopies. The canopy space used for voxelization in this study is indicated by the two horizontal black lines. The figure was created with_BioRender.com. Leaf illustrations by Carolina Levicek. **(b)** Subplot point cloud of a species mixture, and **(c)** the associated occupied voxel per strata with a voxel size of 0.4 m edge length. Red line (lowest crown base height of subplot) to green line (highest tree height of subplot) indicates the canopy space definition used in this study.

### 2.4. Community productivity

For each tree within a subplot, we used diameter at breast height (D) and tree height (H) to calculate aboveground wood productivity over a 5-year period (2012 to 2017). The wood volume (*V_i_*, m³) for each tree *i* was calculated as 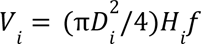 using a form factor *f* of 0.4 in order to approximate the calculated volume of a cylinder to the shape of a tree (Pretzsch, 2009; Kunz et al., 2019).

For each plot *j*, annual wood productivity (AWP, cm³ year^-1^) was calculated as the sum of the annual growth rates of all living trees (*n*) within a plot:

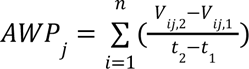

where *V_ij,1_*and *V_ij,2_* are the wood volumes of tree *i* in plot *j* at the beginning (t_1_) and at the end (t_2_) of the study period (2012–2017), respectively.

### 2.5. Leaf trait data and calculation of functional community characteristics

We selected five leaf traits that are linked to the leaf economics spectrum (Wright et al., 2004), and thus capture the resource-use strategies of our tree species: leaf dry matter content (LDMC), specific leaf area (SLA), leaf nitrogen content (LNC), leaf carbon content (LCC), and leaf carbon to nitrogen ratio (C:N). All trait data were obtained from leaves that were collected in the Sardinilla experiment in May and June 2017, i.e. in parallel to the terrestrial laser scanning. In each plot, we collected leaves from up to 11 individuals per species in different height levels along their outer canopy. For further information on leaf trait collection and measurements see Proß et al. (2021). For each subplot, we used plot-specific species mean trait values to calculate a community-weighted mean (CWM) for each of the five leaf traits (LDMC, SLA, LNC, LCC, C:N ratio). Functional diversity (FD) was calculated as the functional dispersion (FDis) of these traits. By using plot-specific trait values, the functional characteristics of our tree communities account for phenotypic trait adjustments in response to varying neighborhood conditions (Davrinche and Haider, 2021), and thus to diversity-induced changes in these traits. As no trait data were available for *T. rosea* in plot A4 (five-species mixture) and *C. odorata* in plot A5 (five-species mixture), we used these species’ mean trait values across all plots to calculate the functional community characteristics of these plots. CWM and FD were computed based on the relative abundance of the species in 2017 using the FD R package (Laliberté and Legendre, 2010).

### 2.6. Statistical analysis

We used generalized linear mixed-effect models (GLMMs) to identify drivers of canopy space filling and community productivity (AWP). To avoid biased inferences associated with logarithmic transformations, we applied a gamma probability distribution and a log-link to ensure homoscedasticity of the residuals (Zuur et al., 2009). Canopy space filling index (CSFI) was modeled as a function of tree species richness, tree density and CWM of leaf dry matter content (CWM-LDMC), leaf nitrogen content (CWM-LNC) and leaf carbon content (CWM-LCC). Moreover, we used the number of living trees within a subplot (i.e. tree density) in 2017 (the year of terrestrial laser scanning and leaf collection) as a further predictor that might affect canopy space filling (Pretzsch, 2014). Given the strong correlation between tree species richness and leaf functional diversity (Pearson correlation *r* = 0.83, *p* < 0.001), we fitted a series of separate models by using functional instead of taxonomic diversity as predictor to assess the importance of both, taxonomic diversity and functional diversity (FD) on CSFI. CWM of SLA and C:N ratio were not considered in the analyses due to their strong correlation with other traits (see Figure 2). All variance inflation factors (VIFs) were <1.8, indicating no collinearity among selected predictors. To test how canopy space filling translates in community productivity and how this relationship changes with tree diversity, we modeled AWP as a function of CSFI and tree diversity (either using FD or tree species richness as predictor). Model selection (backward) was performed using likelihood-ratio tests. All predictors were standardized (mean=0, SD=1) prior to analysis.

**Figure 2.**
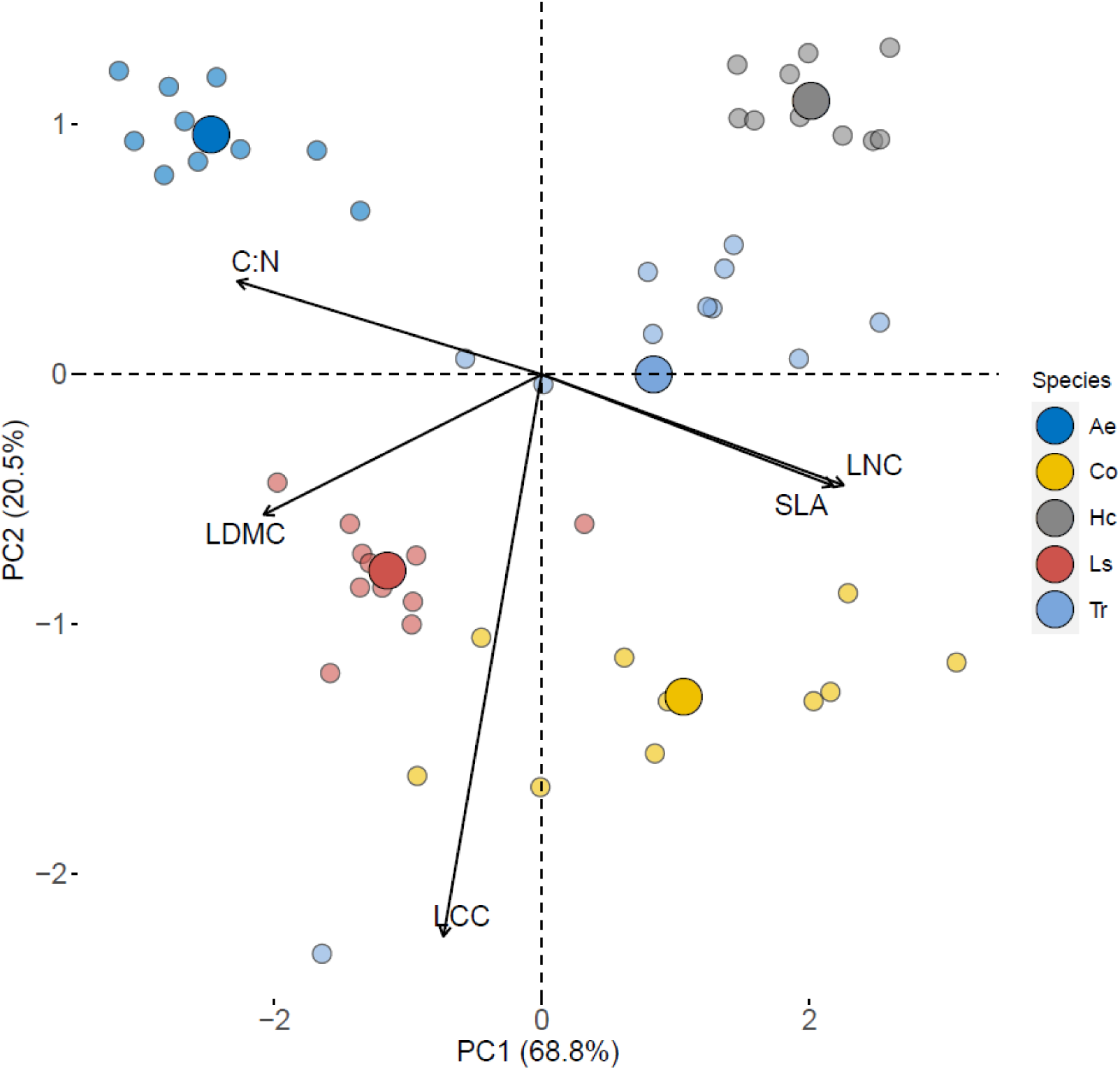
Principal component analysis of five leaf traits. Trait abbreviations: LDMC, leaf dry matter content; SLA, specific leaf area; LNC, leaf nitrogen content; LCC, leaf carbon content; C:N, leaf carbon to nitrogen ratio. Small colored dots represent plot-specific species scores, while large colored dots represent the species’ centroids in the trait space. Species codes: Ae, *Anacardium excelsum*; Co, *Cedrela odorata*; Hc, *Hura crepitans*; Ls, *Luehea seemannii*; Tr, *Tabebuia rosea*.

For all mixed-species communities, we first assessed the bivariate relationships between tree taxonomic and functional (FD and CWM) community characteristics and net biodiversity effects on canopy space filling and AWP by using linear mixed-effects models (LMMs). For each mixture subplot, we calculated net biodiversity effects (NE) as: NE(*x*)*_i_* = x*_obs,i_*– x*_pred,i,_*, where x*_obs,i_* is the observed CSFI or AWP in mixture *i,* and x*_pred,i_* is the sum of CSFI or AWP values measured in the monoculture subplots of the constituent species weighted by their relative abundance (in 2017) in mixture *i*. Positive net biodiversity effects on AWP indicate overyielding (i.e. higher wood volume in a mixture compared to the weighted average of monoculture yields), while negative effects indicate underyielding (i.e., lower wood volume in a mixture compared to the weighted average of monoculture yields).

Second, we used structural equation modeling (SEM) based on LMMs to identify possible pathways by which varying levels of tree species richness in mixtures could change overyielding (i.e. positive net biodiversity effects on AWP). We included direct effects between leaf functional diversity (FD), functional identity (CWM-LNC) and net biodiversity effects on canopy space filling (NE-CSFI) with overyielding. Moreover, we tested for indirect effects of tree species richness, FD and CWM-LNC on overyielding via their effects on NE-CSFI. We hypothesized that positive NE-CSFI are positively related to a higher variation in species with contrasting leaf traits (i.e. higher FD), which would allow communities to fill canopy space more efficiently due to physical niche complementarity (Schmid and Niklaus, 2017). It is also plausible that positive NE-CSFI are the result of a higher dominance of species with more acquisitive leaf traits (i.e. high leaf nitrogen content) within a community, leading to an enhanced canopy space occupation by changing their crown structure (e.g. crown size, crown sinuosity) or leaf area index due to improved photosynthetic rates (Kröber et al., 2014; Pretzsch, 2014). To test this hypothesis, we used CWM-LNC as a proxy for a gradient of acquisitive (high values of LNC) versus conservative resource-use strategies (low values of LNC) of our study species, as CWM-LNC was most strongly correlated with the first axis of a principal component analysis of the five considered leaf traits in our study (Pearson correlation *r* = 0.94, *p* < 0.001;see Figure 2). Lastly, differences among tree species that are not captured by their leaf traits, such as branch traits, could also modulate crown complementarity (Hildebrand et al., 2021) and therefore NE-CSFI. Thus, tree species richness effects on NE-CSFI can play an additional important role in shaping the relationship between NE-CSFI and overyielding. This approach allowed us to explore more mechanistically how taxonomic and functional community characteristics are linked with net biodiversity effects on canopy space filling and overyielding.

For all mixed-effects models (GLMMs and LMMs), plot was used as a random effect to avoid pseudoreplication, and subplots were nested within plots. Importantly, the plot also accounts for the realized compositional differences in tree species among subplots in 90% of the data (in 2 out of 21 plots tree species composition varied between the four subplots due to mortality of one tree species prior to the investigated study period). Model validations were visually assessed and confirmed according to Zuur et al. (2009). All analyses were conducted in R (version 4.3.1; R Core Team, 2023) using the packages factoextra (Kassambara and Mundt, 2020), FD (Laliberté et al., 2014), lme4 (Bates et al., 2015), lmerTest (Kuznetsova et al., 2017), MuMIn (Barton, 2022), piecewiseSEM (Lefcheck, 2016), vegan (Oksanen et al., 2022), tidyverse (Wickham et al., 2019).

## 3. Results

### 3.1. Functional strategies of study species

A principal component analysis (PCA) based on species mean trait values per plot showed that 89.3% of the variation in leaf traits was captured by two dimensions (Figure 2). A substantial amount of variation was explained by acquisitive versus conservative resource-use strategies (PC1: 68.8%). A second orthogonal axis was related to leaf carbon content only (PC2: 20.5%). Consequently, two tree species in our study were associated with conservative (*A. excelsum*, *L. seemannii*) and three species with more acquisitive leaf traits (*T. rosea*, *C. odorata*, *H. crepitans*; Figure S1). *C. odorata* showed the highest and *H. crepitans* the lowest trait variability among plots, which comprise all types of responses to changes in biotic (e.g. tree species richness, tree species composition and tree density) and abiotic (e.g. small-scale variations in water and nutrient supply) plot conditions (Figure 2).

### 3.2. Drivers of canopy space filling

Tree species richness and the community-weighted means of leaf nitrogen content (CWM-LNC) and leaf carbon content (CWM-LCC) explained a substantial amount (76%) of the variation in canopy space filling index (CSFI). CWM of leaf dry matter content (*t* = 0.19, *p* =0.853), tree density (*t* = 1.32, *p* = 0.186) and leaf FD (*t* = 0.05, *p* = 0.133; Figure S2) were not significantly related to CSFI (Figure 3a). We found moderate evidence for an overall increase in CSFI with tree species richness across all levels of tree species richness (monocultures up to five-species mixtures; *t* = 2.03, *p* = 0.042). Although a high variability in CSFI values in monocultures was evident, CSFI was on average 24% higher in five-species mixtures compared to monocultures (Figure 3b). CMW-LNC, however, was the strongest determinant of CSFI (*t* = −6.16, *p* < 0.001; Figure 3a), with communities composed of species with more conservative leaf traits (i.e. lower values of CWM-LNC) exhibiting highest values of CSFI (Figure 3c). CSFI declined with increasing values of CWM-LCC (*t* = −3.96, *p* < 0.001; Figure 3d). These responses were strongly driven by monocultures, indicating a dominant role of species identity effects in explaining changes in canopy space filling across levels of tree species richness (monocultures up to five-species mixtures). For example, leaves of *A. excelsum* had the lowest leaf nitrogen per unit mass on average (Figure S1) and the highest CSFI in monocultures (Figure S3a). The opposite pattern was evident for *C. odorata* and *H. crepitans*.

**Figure 3.**
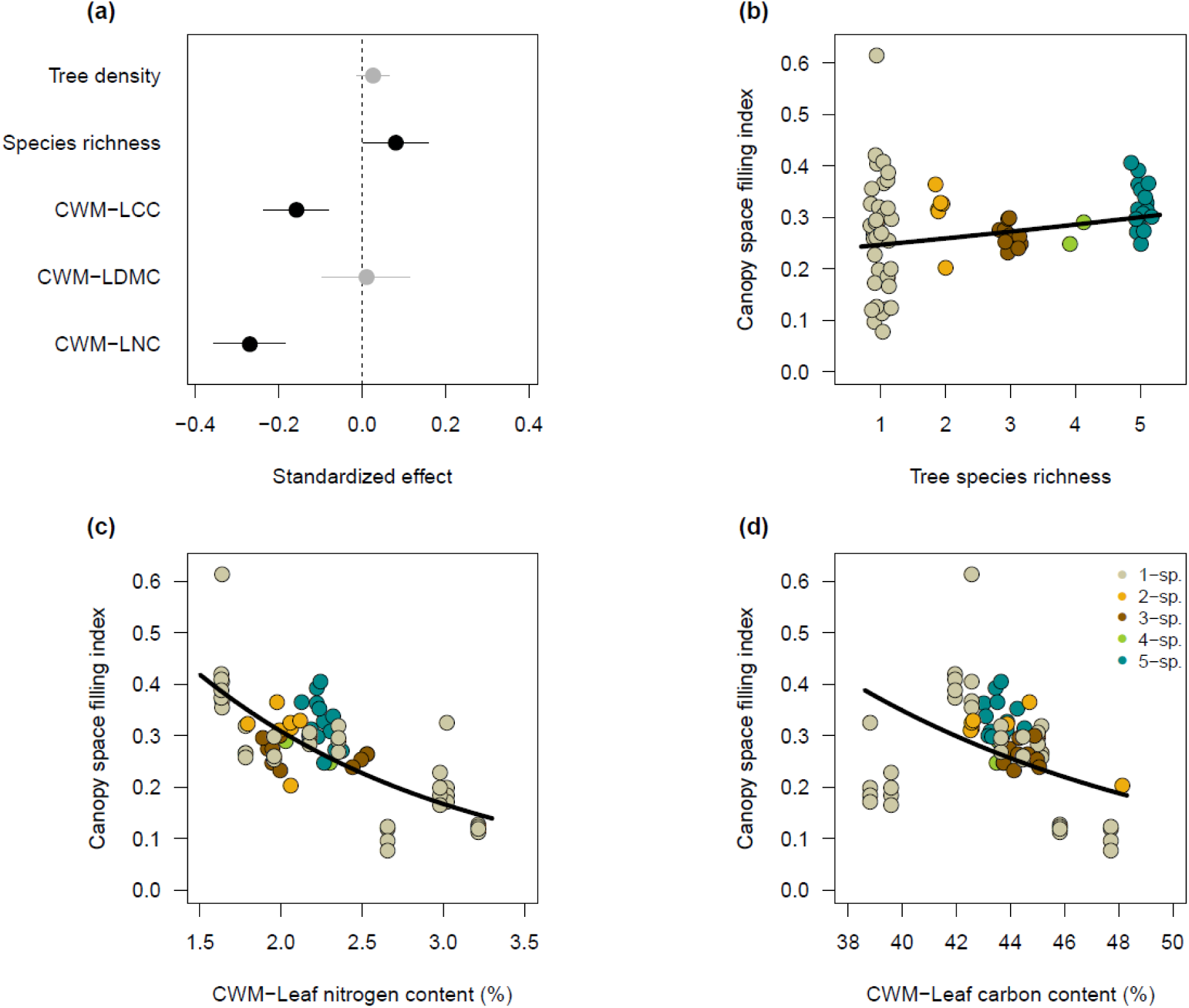
Effects of tree species richness, tree density and community-weighted means (CWMs) of leaf traits (LDMC: leaf dry matter content; LNC: leaf nitrogen content; LCC: leaf carbon content), on canopy space filling index (CSFI). **(a)** Standardized regression coefficients obtained from a generalized mixed-effect model. Points are predicted means, and error bars denote the 95% confidence intervals. Error bars not overlapping with zero indicate significant effects (black points), and vice versa (grey points). **(b)-(d)** Relationship between CSFI and tree species richness, CWMs of leaf nitrogen (CWM-LNC) or leaf carbon content (CWM-LCC). Regression lines correspond to the predicted response of the best-fitted generalized mixed-effect model (see Table S1), while keeping the other covariates constant at their means. Colored dots represent the observed values.

### 3.3. Linking canopy space filling with productivity across the tree species richness gradient

We found strong evidence for an overall increase in annual wood productivity (AWP) with canopy space filling across all levels of tree species richness (*t* = 3.21, *p* = 0.001; Figure 4a and Figure S4a). Communities associated with a high canopy space filling (50% quantile of CSFI; CSFI >0.29) were 61% more productive on average than those with a low canopy space filling (lower than the 50% quantile of CSFI). Notably, AWP in monocultures of conservative species (*A. excelsum, L. seemannii*) was 1.7-times higher on average than those of acquisitive species (*C. odorata*, *H. crepitans*, *T. rosea*; Figure S3b), which coincides with their 76% higher CSFI (Figure S3a). Furthermore, AWP increased with increasing functional dissimilarity of the leaves within the canopy (*t* = 2.43, *p* = 0.015, Figure 4b) and tree species richness (*t* = 2.61, *p* = 0.009; Figure S4b). For example, functionally diverse communities (i.e. higher than the 75% quantile of FD; i.e. FD >8.34) had accumulated 80% more stem volume per year on average than monocultures. Similarly, five-species mixtures were 77% more productive on average compared to monocultures. The canopy space filling-productivity relationship, however, did not vary with leaf FD (interaction term: *t* = 0.83, *p* = 0.404) or tree species richness (interaction term: *t* = 0.56, *p* = 0.559). Canopy space filling index and tree diversity explained 51% (CSFI and FD; Table S2) and 53% (CSFI and tree species richness; Table S3) of the variation in AWP, respectively.

**Figure 4.**
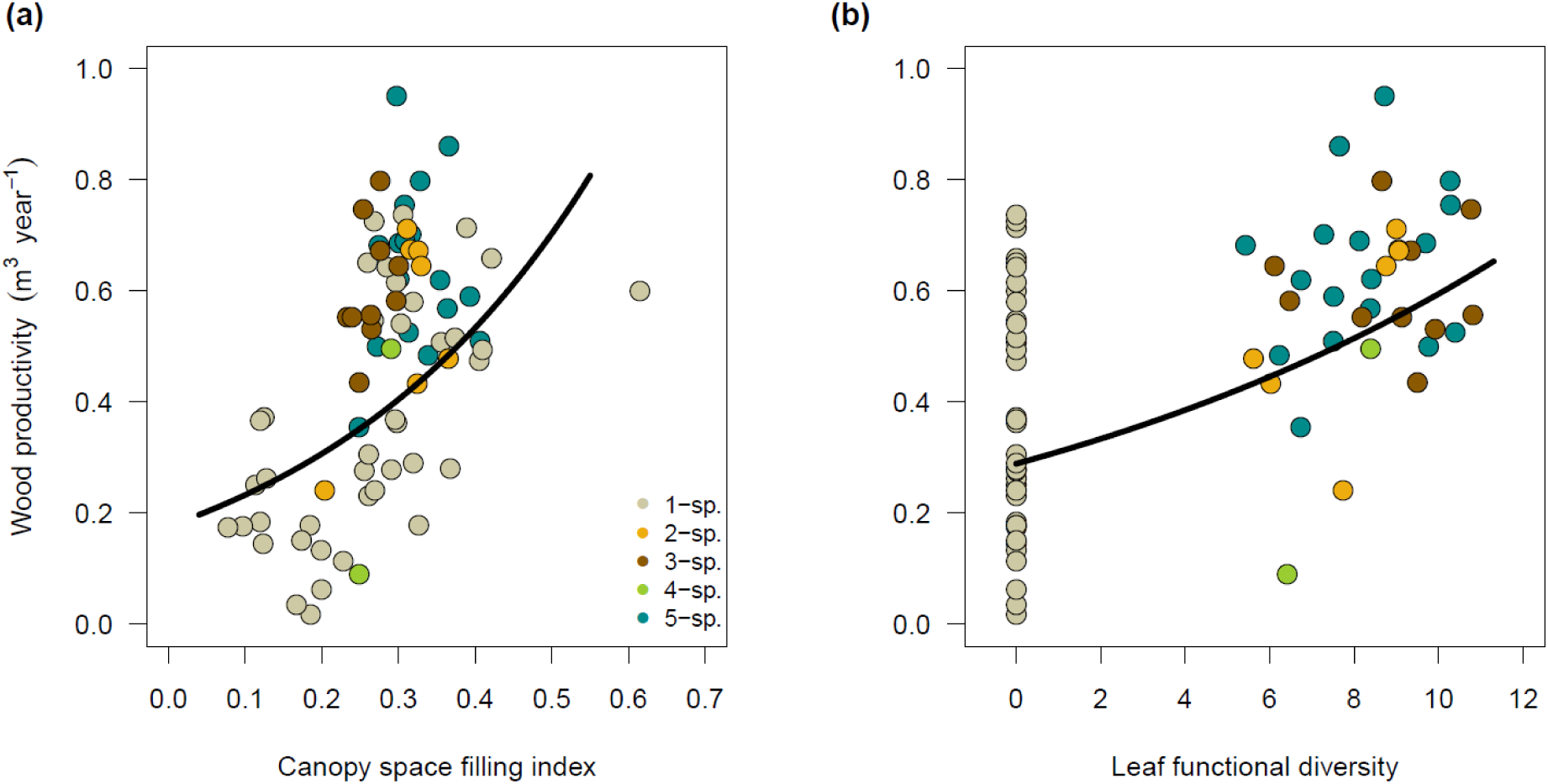
Effects of **(a)** canopy space filling index (CSFI) and **(b)** leaf functional diversity on community productivity (AWP). Regression lines correspond to the predicted response of the best-fitted generalized mixed-effect model (Table S2) while keeping the other covariate constant at its mean. Colored dots represent the observed values.

### 3.4. Relative importance of functional characteristics and diversity-mediated shifts in canopy space filling for overyielding

A large proportion of the mixtures (83%) were more productive than the weighted average productivity of constituent species in monocultures (i.e. most of them overyielded), indicating a strong positive net biodiversity effect on AWP. Moreover, in half (50%) of the mixed-species communities CSFI was greater than expected based on the CSFI values of the same species in monocultures, indicating a moderate positive net biodiversity effect on canopy space filling (NE-CSFI). Leaf FD (*t* = 2.43, *p* =0.025; marginal *R*²: 0.19) and NE-CSFI (*t* = 2.03, *p* =0.054; marginal *R*²: 0.14) were positively associated with changes in overyielding (Figure 5 a,b). On average, overyielding of mixtures that benefited from a tree diversity-induced increase in CSFI (positive NE-CSFI values) was two-fold (99.7%) higher (*t* = 2.51, *p* =0.017; marginal *R*²: 0.15) compared to those with negative NE-CSFI values (Figure 5c) Although NE-CSFI increased with tree species richness (*t* = 2.62, *p* =0.019; marginal *R*²: 0.28), the observed positive relationship between NE-CSFI and overyielding was largely attributable to tree species rich-communities (five-species mixtures). In 76% of the five-species mixtures NE-CSFI was positive, while only 26% of the less species-rich mixtures (two-, three- and four-species mixtures) exhibited positive NE-CSFI values (Figure 5d). Similarly, NE-CSFI increased with increasing CWM-LNC (*t* = 1.97, *p* =0.059; marginal *R*²: 0.14; Figure 5e).Moreover, NE-CSFI tended to increase with leaf FD, but this effect was weak and not significant (*t* = 1.22, *p* =0.232; marginal *R*²: 0.04; Figure 5f).

**Figure 5.**
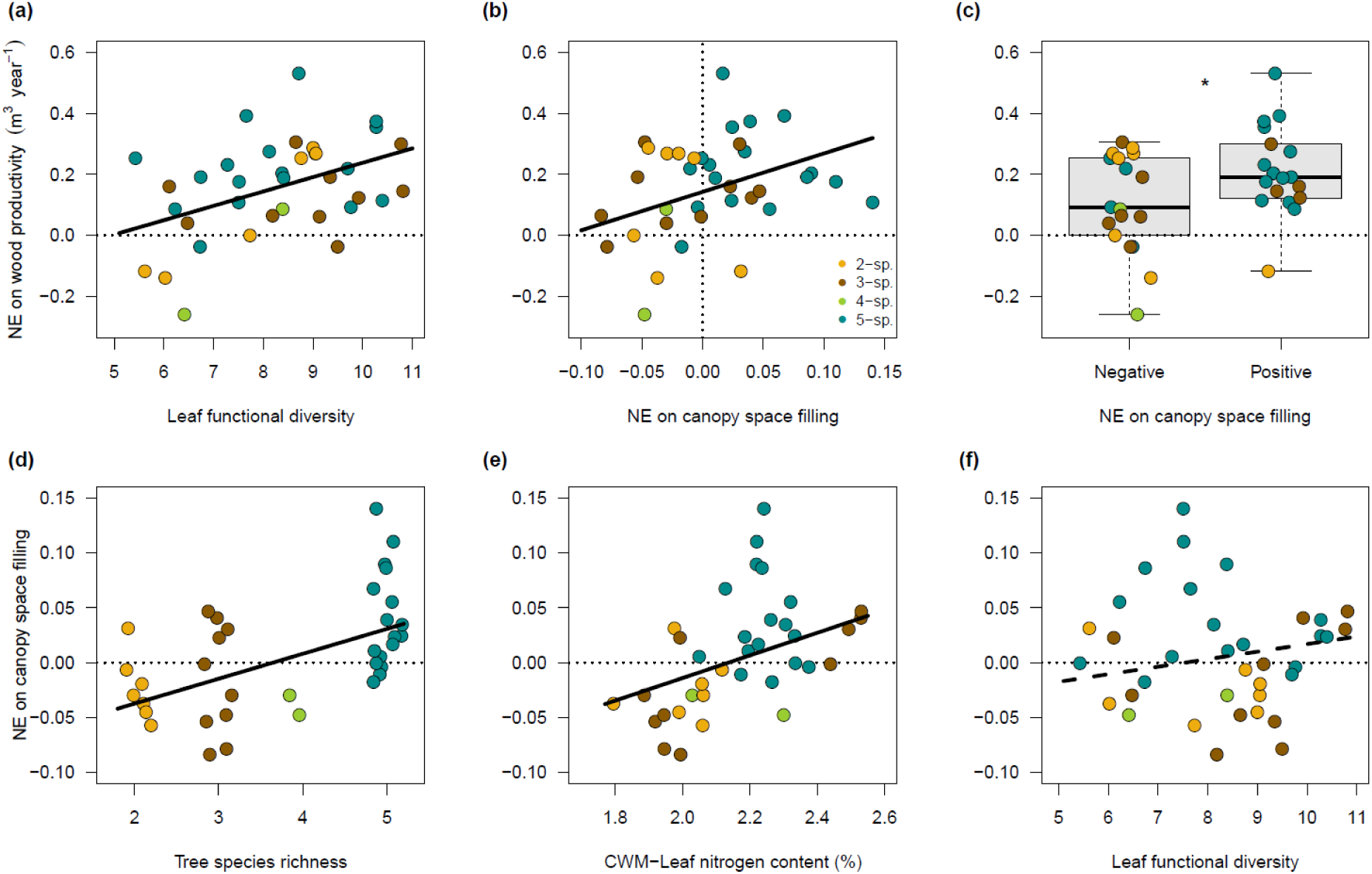
Bivariate relationships between overyielding (NE on wood productivity) and **(a)** leaf functional diversity and **(b)** net biodiversity effects on canopy space filling. **(c)** Changes in overyielding with positive and negative net biodiversity effects (NE) on canopy space filling. Bivariate relationships between net biodiversity effects on canopy space filling and **(d)** functional diversity and **(e)** tree species richness. Regression lines are linear mixed-effect model fits and colored dots represent the observed values. The comparison in the differences of means in panel **(c)** is also based on a linear mixed-effects model; * *p* < 0.05.

Structural equation modeling (SEM) indicated that net biodiversity effects on canopy space filling (standardized path coefficient: 0.483, *p* = 0.024) and leaf FD (standardized path coefficient: 0.480, *p* = 0.016) were of equal importance for explaining overyielding (Figure 6). However, we found no support for our hypotheses that a higher functional dissimilarity in leaf traits within a canopy (leaf FD: standardized path coefficient: 0.136, *p* = 0.387) or a higher abundance of acquisitive species within a community (CWM-LNC: standardized path coefficient: 0.233, *p* = 0.233) would lead to diversity-enhanced canopy space filling (i.e. higher NE-CSFI), which in turn would translate into higher overyielding. Tree species richness had no significant effect on the functional community characteristics, as neither leaf FD (standardized path coefficient: 0.167, *p* = 0.517) nor CWM-LNC (standardized path coefficient: 0.304, *p* = 0.156) were affected by tree species richness. Instead, tree species richness promoted overyielding indirectly, via increasing NE-CSFI (standardized path coefficient: 0.450, *p* = 0.056). CWM-LNC was negatively, but not significantly related to overyielding (standardized path coefficient: 0.195, *p* = −0.295), indicating that communities composed of more conservative species tended to have higher overyielding.

**Figure 6.**
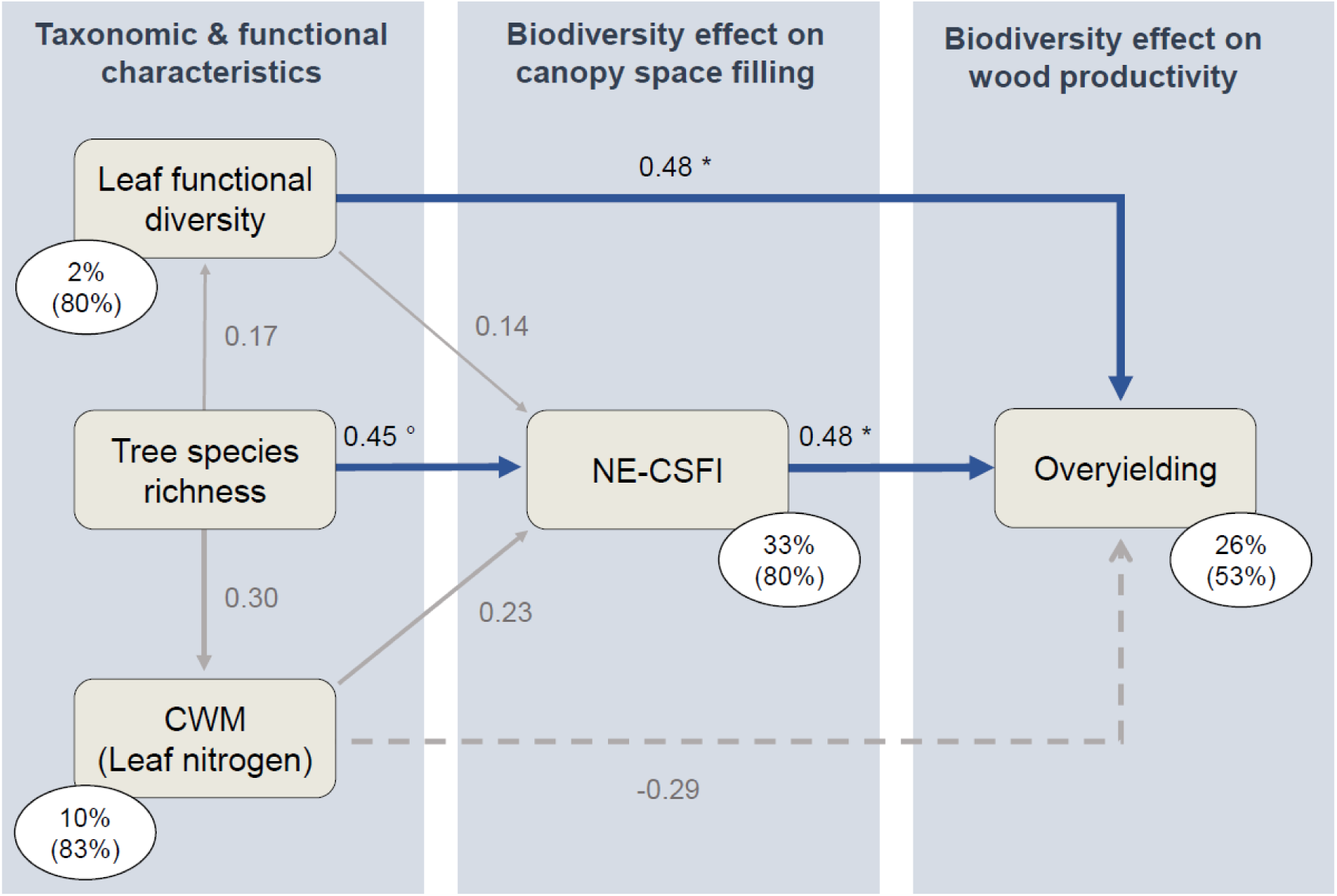
Structural equation model linking taxonomic (tree species richness) and functional (leaf functional diversity and community-weighted mean of leaf nitrogen content) community characteristics with net biodiversity effects on canopy space filling (CSFI) and wood productivity (overyielding). The blue arrows indicate significant (* *p* < 0.05) and marginal significant (° *p* < 0.10) positive pathways. Solid and dotted gray arrows indicate nonsignificant (*p* > 0.10) positive (solid lines) and negative (dotted lines) pathways, respectively. Numbers next to arrows are standardized path coefficients, and arrow width is proportional to standardized path coefficients. The explained variance by fixed effects is given as percentage values; the variance explained by fixed and random effects are in parentheses. The model fitted the data well, as indicated by Fisher’s *C* = 3.703, *p* = 0.448, d.f. = 4. CWM: community-weighted mean.

## 4. Discussion

Complementarity among trees in the canopy space is considered as a crucial ecological mechanism underlying positive biodiversity-productivity relationships in forests. Using a voxel grid approach based on point cloud data from terrestrial laser scans in a controlled tree diversity experiment, we found that species identity effects were more important in driving canopy space filling than taxonomic or functional tree diversity. In contrast, tree diversity and canopy space filling together explained community productivity, with communities associated with densely-packed and functionally diverse canopies being the most productive ones. Our results provide further evidence that net biodiversity effects on wood productivity can be attributed to enhanced canopy space filling and leaf functional diversity.

### 4.1. Drivers of canopy space filling

We found that the mixed-species communities had higher canopy space filling than the monocultures. This is consistent with several other studies (Pretzsch, 2014; Jucker et al., 2015; Georgi et al., 2022; Martin-Blangy et al., 2023), but contrasting patterns have also been observed (Seidel et al., 2013). Important aspects of such studies are the dominance structure in the tree communities and the length of the diversity gradient. In observational studies, one or a few tree species often have a higher mixing proportion than the other species and can dominate the result (Seidel et al., 2013; Georgi et al., 2022). However, there has been a lack of studies investigating diversity effects on canopy space filling under experimentally controlled conditions across a longer tree species richness gradient. Due to the differences in growth rates of the fast-growing, light-intermediate species and slow-growing shade-tolerant species planted in Sardinilla (see methods), vertical stratification (i.e. physical complementarity among trees in the use of vertical space) is likely an important mechanism for the observed mixing effects on canopy space filling (Ray et al., 2023; Schmid and Niklaus, 2017). However, the second potential mechanism, diversity-induced crown plasticity, is also a driver of crown complementarity here, as reported not only from temperate (Pretzsch, 2014; Georgi et al., 2021) and subtropical (Kunz et al., 2019; Hildebrand et al., 2021) forests, but also from the tropical Sardinilla experiment (Guillemot et al., 2020). In combination, plasticity-mediated changes in crowns lead to canopy space filling through investing in space-foraging branches and investing in the trunk, which results in a vertical distribution of crown volume. Crowns become more dense in volume and more sinuous in mixtures (Guillemot et al., 2020).

Conflicting results from other studies may also be due to differences in the technical approach to quantifying canopy space filling and in the definitions of the canopy space. The great advantages of terrestrial laser scanning (TLS) for measuring the complex structures of forests are now widely recognized (Lines et al., 2022). TLS offers a much wider range of assessment options and helps to avoid errors and inaccuracies. Notable among these are the elimination of the need to use standard geometric shapes to quantify individual tree crowns, and the great flexibility in the exact delineation of the crown space. Regarding the latter, Georgi et al. (2022) showed that the different definitions of the canopy space can even lead to contradictory results in the analysis of mixing effects on canopy space filling. For our study, however, it is important to note that several of the aspects outlined by Georgi et al. (2022) are only relevant when studying mature forests in an observational setting. A wide range of tree sizes (and thus generally heterogeneous age distribution) means that the canopy can be subdivided into different layers: overstory, midstory (“canopy”), understory, and shrub layer. In such forests, it is crucial how exactly the canopy space is defined and which tree sizes are included in the analyses (Georgi et al., 2022). In our experimental study site, however, all trees were of the same age (Potvin and Dutilleul, 2009; Scherer-Lorenzen et al., 2005) and were still relatively young at the time of data collection (16 years).

While the relationship between tree species richness and canopy space filling was moderately strong, we found a very strong relationship between the community-weighted means of leaf nitrogen and leaf carbon contents and canopy space filling. This shows that in addition to confirming the niche complementarity hypothesis, the mass-ratio hypothesis was also supported. It is important to note that the two hypotheses are not necessarily mutually exclusive (Mokany et al., 2008; Chiang et al., 2016). Communities with a high abundance of species with conservative leaf traits, particularly *A. excelsum*, but also *L. seemannii*, were those with most densely packed canopies. In contrast, the two acquisitive tree species *H. crepitans* and especially *C. odorata* showed low canopy space filling. The mixture of tree species with different strategies and architectural traits thus leads to a high canopy space filling even in relatively young plantations, which has also been shown in terms of stand structural complexity (Perles-Garcia et al., 2022; Ray et al., 2023).

### 4.2. Canopy space filling and wood productivity

We found a strong increase in wood productivity with increasing canopy space filling. This is theoretically expected, as higher canopy space filling is thought to lead to higher light absorption, and thus, higher biomass productivity. However, this has only been partially confirmed in studies, and it has been repeatedly stated that this may be due to insufficient recording of the actual filling of the entire canopy space (Juchheim et al., 2017; Martin-Blangy et al., 2023). The complete capture of canopy space filling by TLS under the controlled conditions of a forest BEF experiment allowed us to rigorously test this assumption, and our results provide strong evidence that crown complementarity, underlying the increased canopy space filling, is an important mechanism for enhanced forest productivity. By analyzing canopy space filling at the subplot level, we can confirm what Guillemot et al. (2020) suggested based on the modeling of individual-tree point clouds: namely, that changes in canopy space filling are more important than crown volume for overyielding in this experiment. This coincides with another previous finding from Sardinilla, where intraspecific plastic changes in crown shape have been identified as a mechanism for higher light capture and productivity, respectively, in mixed-species communities (Sapijanskas et al., 2014).

### 4.3. Exploring processes underlying overyielding in mixed-species tree communities

We found that overyielding arose via two equally important processes. First, via higher leaf functional diversity and second, via positive net biodiversity effects on canopy space filling. Intra- and interspecific variation in light use efficiency of constituent species has been shown to generate a higher light use efficiency of mixtures, which in turn translates into overyielding (Williams et al., 2021). Given that our leaf trait data accounted for diversity-induced changes within and among tree species, a higher diversity in leaf traits may allow mixed-species communities to exploit available light resources more efficiently (Plekhanova et al., 2021), leading to reduced competition for light and higher productivity, respectively. Moreover, it is conceivable that community-level light interception increases with increasing canopy leaf diversity due to niche complementarity (Sapijanskas et al., 2014; Niklaus et al., 2017), resulting in enhanced overyielding.

Positive net biodiversity effects on canopy space filling can emerge from multiple processes, such as inter- and intra-specific differences in tree growth strategies and resulting vertical stratification as well as from variation in inner and outer crown morphology and crown architecture, in addition to the effect of leaf functional diversity (Kunz et al., 2019; LaRue et al., 2023; Pretzsch, 2014; Jucker et al., 2015; Hildebrand et al., 2021). Shade tolerance is a crucial trait in ecology that affects interactions among plants because of its link to those functional traits that determine a tree’s ability to compete for light (Valladares and Niinemets, 2008; Kunstler et al., 2016). For example, fast-growing species are often associated with a low and slow-growing species with a high shade tolerance (Sendall et al., 2016). Thus, communities composed of species with a high variation in shade tolerance can have more vertically structured canopies (Morin et al, 2011; Ray et al., 2023), as shade-intolerant species tend to dominate the upper canopy layers, while shade-tolerant species, which are comparatively shorter, tend to occupy the lower layers of the canopy (Niinemets, 2010). This might be the reason that *L. seemannii* and *A. excelsum* have rounded dense crowns, while both have deeper crowns and higher leaf area index (Kitajima et al., 2005). Our results show that positive net biodiversity effects on canopy space filling predominantly arose in species-rich communities (i.e. five-species mixtures), while this effect was less distinct in less species-rich communities (i.. e. two- and three-species mixtures). This might be attributable to the fact that in species-rich communities of our experiment, interspecific differences in shade tolerance and growth rates already resulted in vertical stratification (Schnabel et al., 2019), which likely contributed to the higher canopy space filling we observe here. Several studies have shown that an increase in forest productivity result from differences in shade tolerance, which in turn translates into greater crown complementarity for light interception (Morin et al., 2011; Toigo et al., 2018; Zhang et al., 2012; Searle and Chen, 2020; Ray et al., 2023). Thus, trees can alter their traits towards more acquisitive values to reach the upper canopy faster (Pellis, 2004). This is in line with our finding that NE-CSFI shifted from negative to positive with increasing CWM-LNC, meaning when communities are dominated by species with acquisitive leaf traits (i.e. high leaf nitrogen content). However, this process was not evident considering multiple pathways simultaneously in the SEM. Morin et al. (2011) also found that mixed forests have a higher leaf area index compared to monospecific stands due to increased canopy space filling. The vertical canopy extension or canopy closure in mixed forests allows for the arrangement of more foliage layers from the canopy top to base, resulting in higher light interception in mixed-species stands (Williams et al., 2021).

The positive relationship between NE-CSFI and NE-productivity could also be attributed to branching characteristics, such as the intensity and accumulation of biomass in branches. These features can alter the plasticity of the inner crown and, in some cases, compensate for outer crown plasticity when functional diversity is low. For example, heterospecific neighbors that are less plastic with respect to their outer crown structure, benefit from an increasing divergence in their branching intensity, which in turn facilitates the spatial adaptation of their branches in the canopy, resulting in enhanced crown complementarity (Hildebrand et al., 2021). Furthermore, the total number of branches formed per branch, known as the ‘bifurcation ratio,’ tends to increase with an increase in light availability and foliage aggregation is influenced by the frequency of branching (Niinemets, 2010). Additionally, tree diversity modulates biomass allocation in branches, which favors a higher variation in crown size and inner crown properties of differing crown shapes among species (Kunz et al., 2019; Guillemot et al., 2020). This suggests that phenotypic changes in branch traits are a further important process that lead to greater complementarity in light-use strategies and canopy space filling, respectively. All these processes, however, are expected to become stronger in species-rich mixtures, which explains why positive net biodiversity effects on canopy space filling were driven by five-species mixtures in our study.

Overall, our study emphasizes the importance of tree species selection in restoration projects that aim at maximizing carbon capture. Tree species with contrasting resource-use strategies allow communities to occupy canopy space more efficiently, and thereby optimizing light capture and enhancing productivity. Consequently, promoting functionally diverse canopies seems to be a promising strategy in forest restoration to meet climate change objectives.

## Acknowledgments

We are grateful to the people who were dedicated to managing the Sardinilla experiment over the years, in particular to José Monteza, Lady Mancilla, and their fieldworkers. We are also grateful to Karl Friedrich Reich and Carsten Hess for their support in TLS data acquisition.

## Author contributions

The Sardinilla experiment was designed and implemented by CP. GvO, AF, MK and HB conceived the idea of the study. MK collected the TLS data. TP and SH collected the leaf data and conducted trait measurements. MH, TR, MK and LG compiled the TLS data. TR and FS compiled the tree growth data. TR and AF analyzed the data and created the figures with assistance from PMB. TR wrote the first draft of the manuscript with assistance from AF and GvO. All authors contributed substantially to revisions.

## Supplementary Materials for

**Table S1.** Results of the best-fitted generalized linear mixed-effects model for canopy space filling index (CSFI). The variance explained by the fixed effects alone (marginal *R*^2^) and by both the fixed and random effects (conditional *R*^2^). Estimates were standardized (mean=0, SD=1). CWM-LNC: leaf nitrogen content, CWM-LCC: leaf carbon content, SR. tree species richness, SE: standard error, SD: standard deviation.

|  | Estimate | SE | *t*-value | *p*-value |
| --- | --- | --- | --- | --- |
| Intercept | -1.334 | 0.043 | -31.365 | < 0.001 |
| CWM-LNC | -0.270 | 0.044 | -6.163 | < 0.001 |
| CWM-LCC | -0.158 | 0.040 | -3.962 | < 0.001 |
| SR | 0.081 | 0.040 | 2.032 | 0.042 |
| SD (plot) | 0.105 |  |  |  |
| SD (residuals) | 0.137 |  |  |  |
| $R^2$ (marginal) | 0.850 | | | |
| $R^2$ (conditional) | 0.860 | | | |

**Table S2.** Results of the best-fitted generalized linear mixed-effects model for community productivity using leaf functional diversity (FD) as a measure of tree diversity. The variance explained by the fixed effects alone (marginal *R*^2^) and by both the fixed and random effects (conditional *R*^2^). Estimates were standardized (mean=0, SD=1). CSFI: canopy space filling index, SE: standard error, SD: standard deviation.

|  | Estimate | SE | *t*-value | *p*-value |
| --- | --- | --- | --- | --- |
| Intercept | -0.959 | 0.133 | -7.223 | < 0.001 |
| CSFI | 0.244 | 0.076 | 3.209 | 0.001 |
| FD | 0.310 | 0.128 | 2.428 | 0.015 |
| SD (plot) | 0.304 |  |  |  |
| SD (residuals) | 0.292 |  |  |  |
| $R^2$ (marginal) | 0.511 | | | |
| $R^2$ (conditional) | 0.766 | | | |

**Table S3.** Results of the best-fitted generalized linear mixed-effects model for community productivity using tree species richness (SR) as a measure of tree diversity. The variance explained by the fixed effects alone (marginal *R*^2^) and by both the fixed and random effects (conditional *R*^2^). Estimates were standardized (mean=0, SD=1). CSFI: canopy space filling index, SE: standard error, SD: standard deviation.

|  | Estimate | SE | *t*-value | *p*-value |
| --- | --- | --- | --- | --- |
| Intercept | -0.980 | 0.143 | -6.868 | < 0.001 |
| CSFI | 0.234 | 0.075 | 3.128 | 0.002 |
| SR | 0.337 | 0.129 | 2.611 | 0.009 |
| SD (plot) | 0.322 |  |  |  |
| SD (residuals) | 0.290 |  |  |  |
| $R^2$ (marginal) | 0.525 | | | |
| $R^2$ (conditional) | 0.787 | | | |

**Figure S1.**
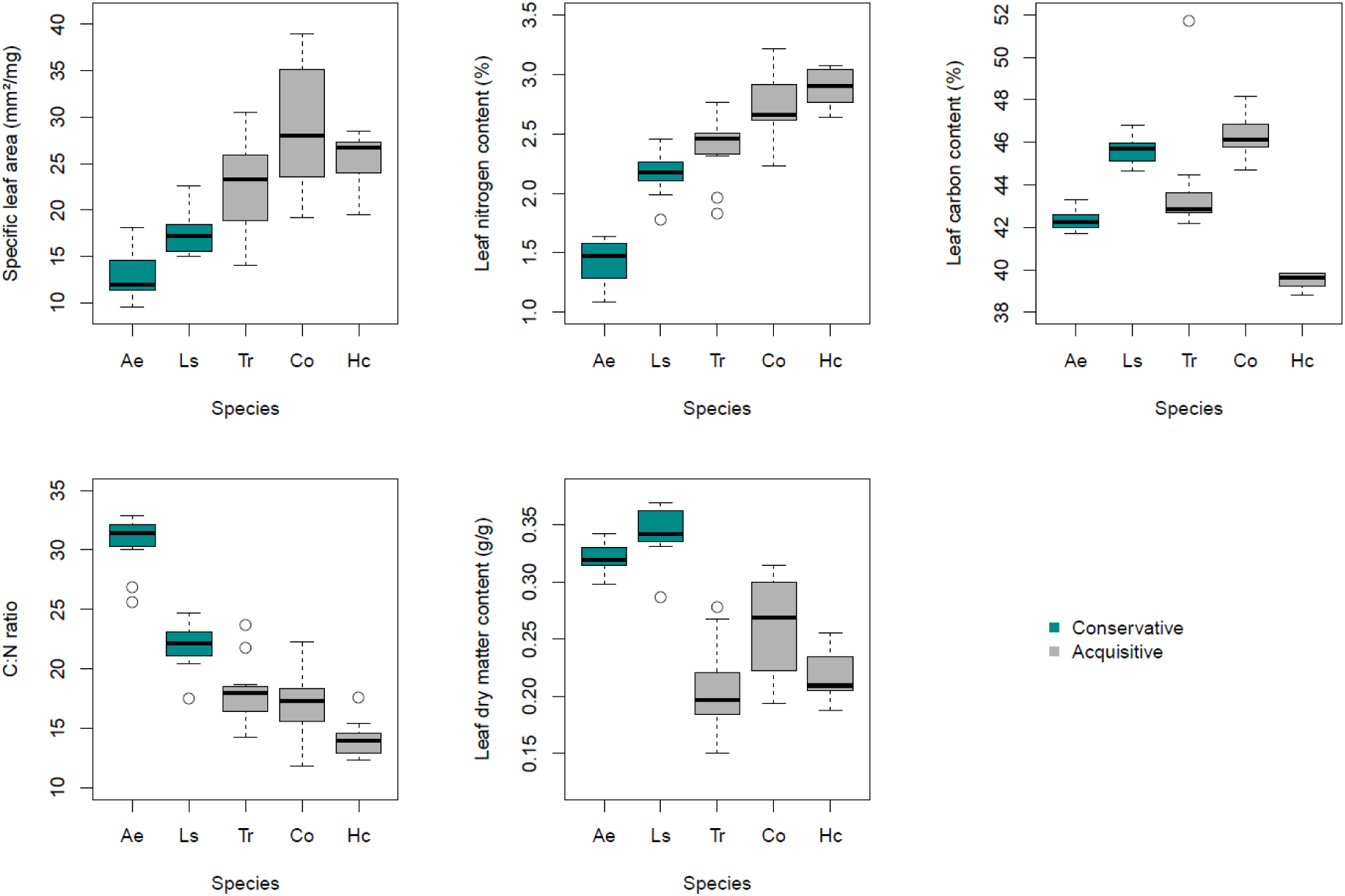
Changes in leaf traits (specific leaf area, leaf nitrogen content, leaf carbon content, carbon nitrogen ratio and leaf dry matter) with tree species. Species are classified as those with conservative and acquisitive resource-use strategies based on the results of a PCA (see Figure 2). Boxplots show the median (horizontal black lines), the 25 and 75% percentiles (edges of the box), and 1.5 times the interquartile range (whiskers) of observed leaf traits. Open circles indicate trait values that are greater or smaller than 1.5 times the interquartile range.

**Figure S2.**
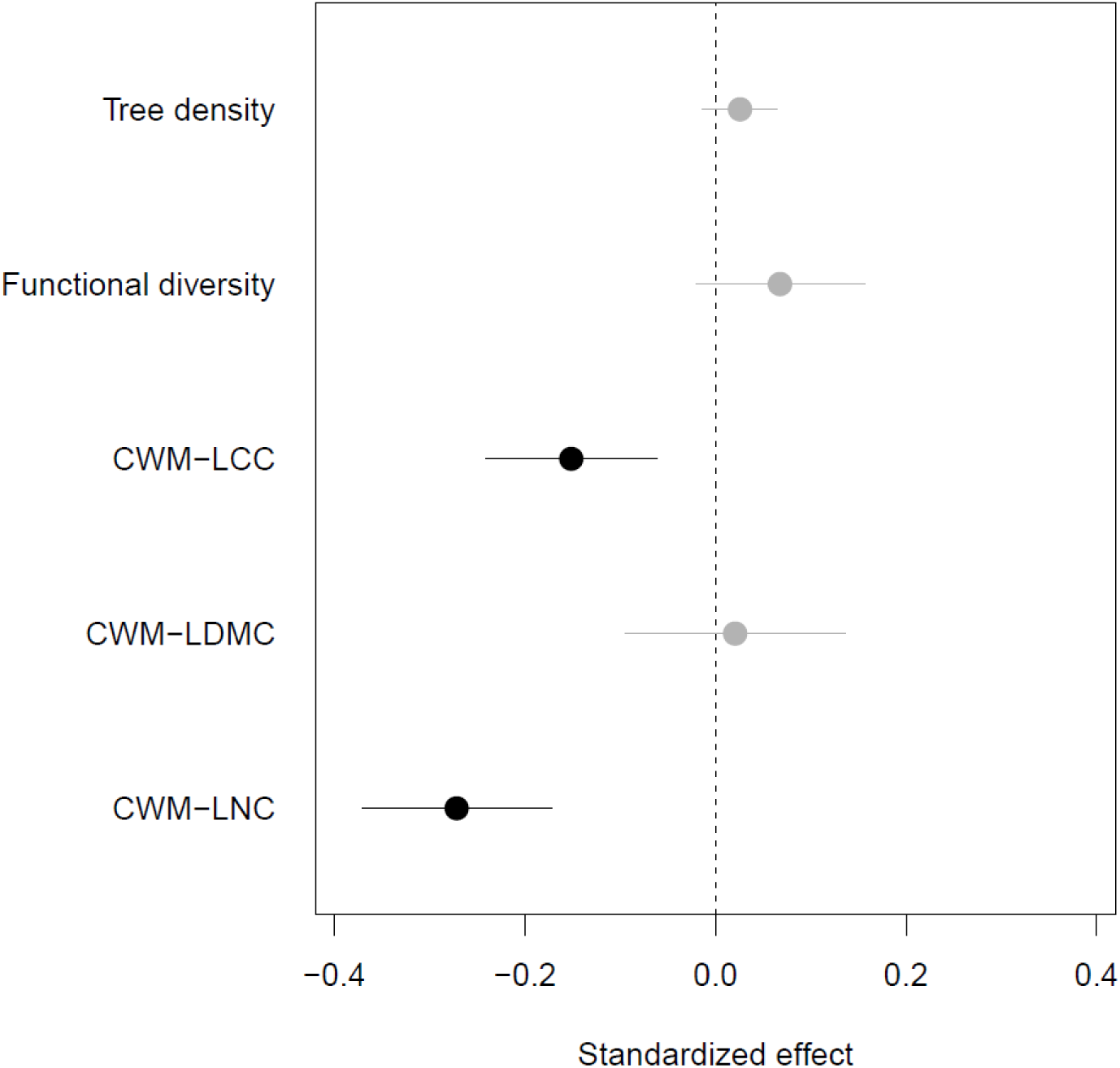
Standardized regression coefficients obtained from a generalized mixed-effect model using functional diversity of leaf traits. Points are predicted means, and error bars denote the 95% confidence intervals. Points are predicted means, and error bars denote the 95% confidence intervals. Error bars not overlapping with zero indicate significant effects (black points), and vice versa (gray points). CWM: community-weighted mean of a leaf trait. LDMC, leaf dry matter content; LNC, leaf nitrogen content; LCC, leaf carbon content.

**Figure S3.**
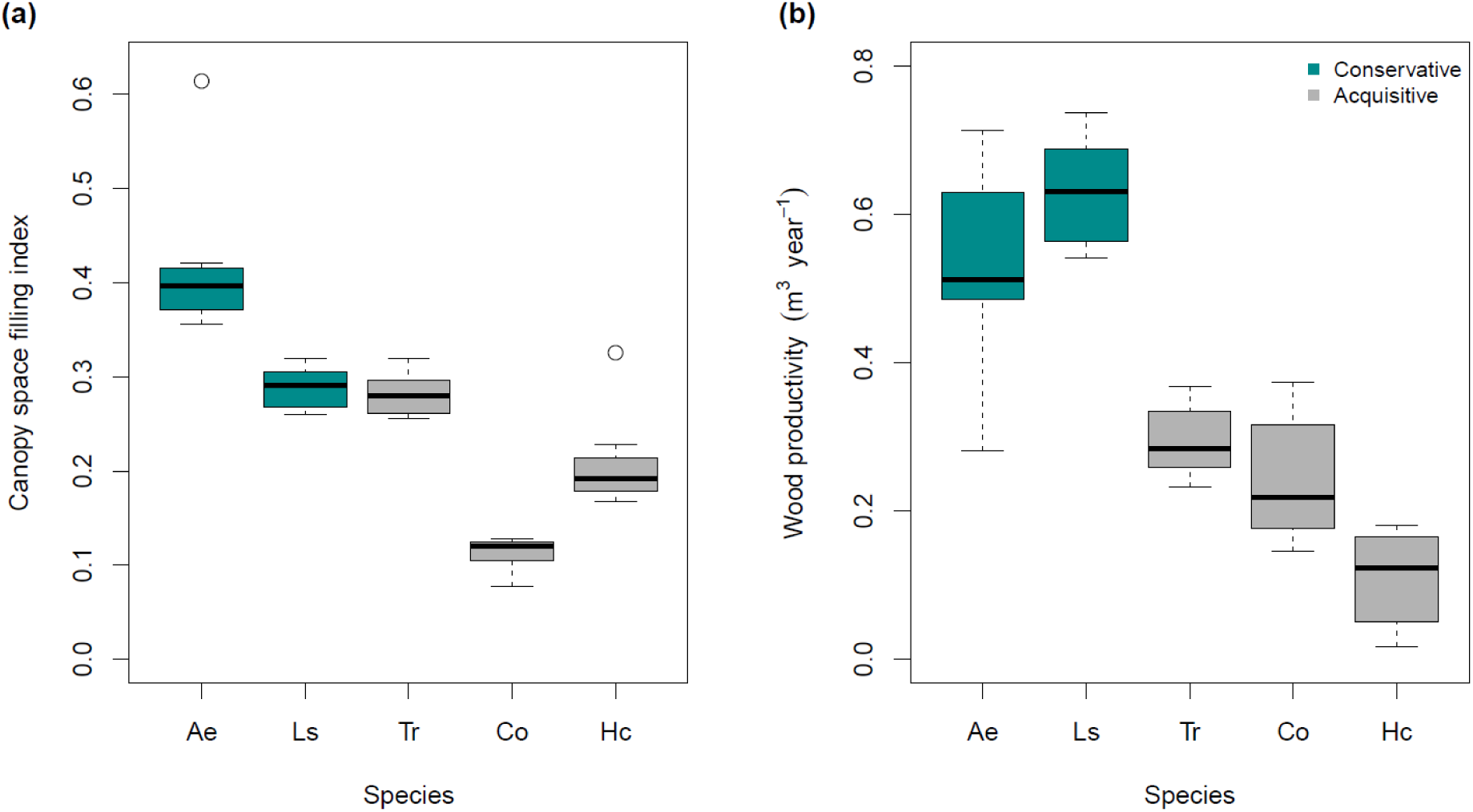
(a) Changes in canopy space filling index (CSFI) and (b) community productivity with tree species in monocultures. Species are classified as those with conservative and acquisitive resource-use strategies based on the results of a PCA (see Figure 2). Boxplots show the median (horizontal black lines), the 25 and 75% percentiles (edges of the box), and 1.5 times the interquartile range (whiskers) of observed CSFI..Open circles indicate CSFI values that are greater or smaller than 1.5 times the interquartile range.

**Figure S4.**
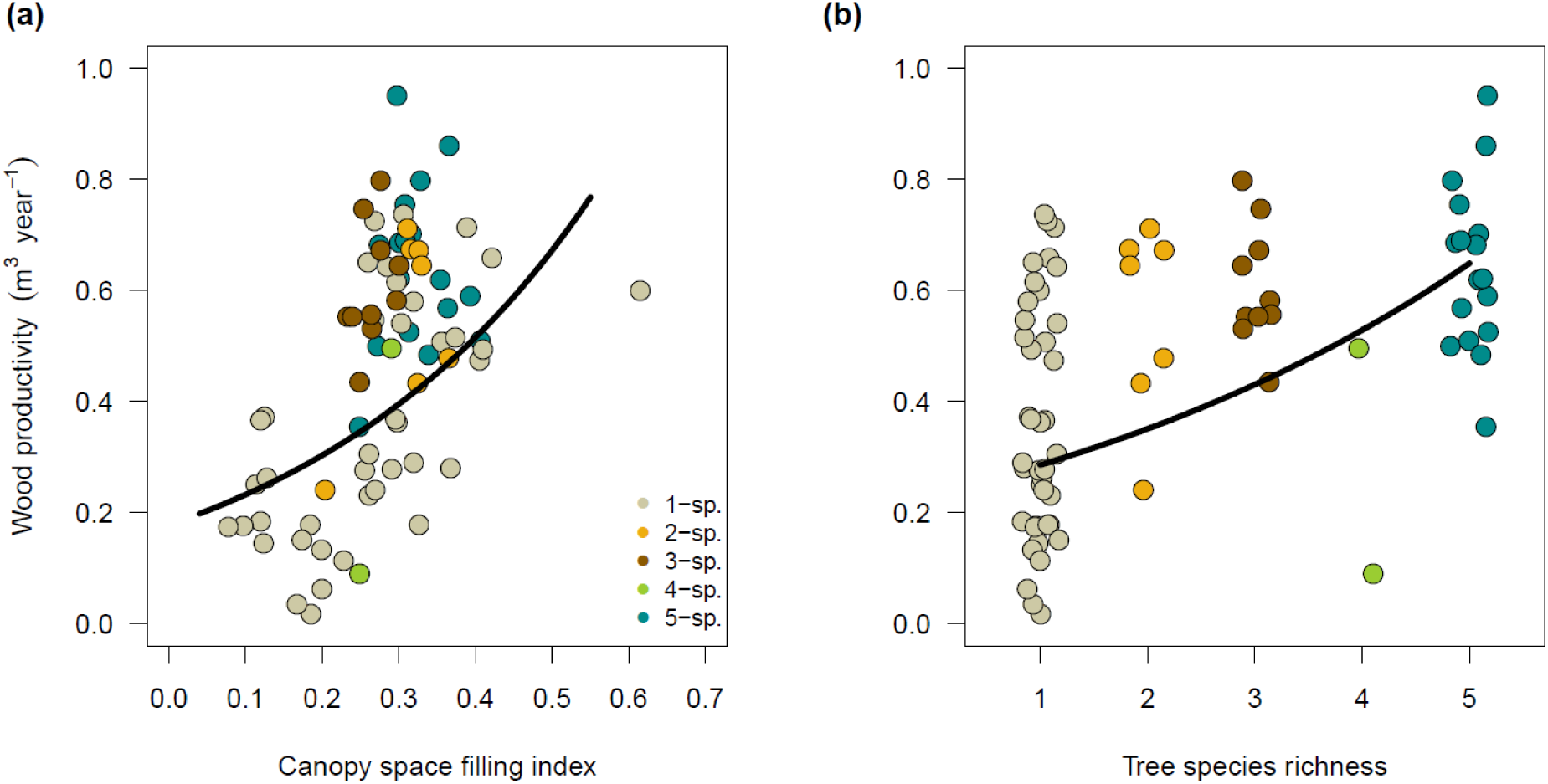
Effects of **(a)** canopy space filling index (CSFI) and **(b)** tree species richness on community productivity. Regression lines correspond to the predicted response of the best-fitted generalized mixed-effect model (Table S3) while keeping the other covariate constant at its mean. Colored dots represent the observed values.

